# Differentiation of vaginal cells from epidermal cells using morphological and autofluorescence properties: Implications for sexual assault casework involving digital penetration

**DOI:** 10.1101/2023.03.30.534941

**Authors:** Sarah Ingram, Arianna DeCorte, Amanda Gentry, M. Katherine Philpott, Taylor Moldenhauer, Sonja Stadler, Cory Steinberg, Jonathan Millman, Christopher J. Ehrhardt

## Abstract

Analysis of DNA mixtures from sexual assault evidence is an ongoing challenge for DNA casework laboratories. There is a significant need for new techniques that can provide information as to the source of DNA, particularly for sexual assault samples that do not involve semen. The goal of this study was to develop a new biological signature system that provides additional probative value to samples comprised of mixtures of epidermal and vaginal cells, as may be observed in cases involving digital penetration. Signatures were based on morphological and autofluorescence properties of individual cells collected through Imaging Flow Cytometry (IFC). Comparisons to reference cell populations from vaginal tissue and epidermal cells collected from hands showed strong multivariate differences across >80 cellular measurements. These differences were used to build a predictive framework for classifying unknown cell populations as originating from epithelial cells associated with digital penetration or epidermal tissue. As part of the classification scheme, posterior probabilities of specific tissue group membership were calculated for each cell, along with multivariate similarity to that tissue type. We tested this approach on cell populations from reference tissue as well as mock casework samples involving digital penetration. Many more cells classifying as non-epidermal tissue were detected in digital penetration samples than control hand swabbings. Minimum interpretation thresholds were developed to minimize false positives; these thresholds were also effective when screening licked hands, indicating the potential utility of this method for a variety of biological mixture types and depositional events relevant to forensic casework. Results showed that samples collected subsequent to digital penetration possessed markedly higher numbers of cells classifying as vaginal tissue as well as higher posterior probabilities for vaginal tissue (≥ 0.90) compared to cell populations collected from hands without prior contact with vaginal tissue. Additionally, digital penetration cell populations may be resolved from saliva cell populations and other non-target tissue types.

## Introduction

Serological testing of sexual assault evidence has long been used by forensic casework units for both presumptive and confirmatory identification of certain biological fluids and cell types. The majority of methods are based on either chemical reactions with body fluid components (e.g., acid phosphatase for seminal fluid, amylase for saliva) or direct observation of cells (e.g., microscopy for sperm cells) and have well-known limitations in terms of sensitivity, specificity, resource allocation, and/or consumption of sample [1]. Moreover, there are cell types for which no validated presumptive or confirmatory test exist such as epidermal cells or vaginal cells. Consequently, for mixture samples that primarily involve these cell types—as would be expected from samples resulting from digital penetration of the vaginal cavity—it may be impossible to infer source tissue. In court this can lead to multiple competing propositions for the presence of non-self, female DNA from a swab of an alleged perpetrator’s fingers following digital vaginal penetration. For example, the defendant may argue that the victim’s DNA may have been transferred to their hands through touch/contact (e.g., handshake) or from deposition of victim’s saliva that was unrelated to the crime. Laboratory techniques that can discriminate between these scenarios can therefore help to reduce the range of possibilities that might be considered to explain the DNA results in these instances.

One possible approach to address this gap is to harness the combinatorial discriminating power of multiple detectable cellular features within a biological sample. Previous studies have shown that morphological and autofluorescence profiles collected through Imaging Flow Cytometry (IFC) can be used to differentiate reference epithelial cell populations originating from vaginal, epidermal, and buccal tissue [2,3]. Although promising, these signatures have not been tested on mixture samples approximating those encountered in casework. Therefore, the goal of this study was to examine the robustness of morphological and autofluorescence signatures for detecting vaginal and epidermal cells in samples collected subsequent to intimate contact. This mixture system was initially chosen due to the paucity of serological techniques for detecting either cell type and its potential applications for sexual assault casework involving digital penetration. Cell type signatures were initially characterized in reference samples from vaginal and epidermal tissue. To further enhance the relevance for forensic casework, efforts were made to identify a reliable saliva signature, given saliva is often an alternative explanation for the presence of DNA in sexual assault cases.

Multivariate differences identified between IFC measurements on these various tissue types were used to construct prediction algorithms for classifying source cell populations from an unknown or blinded sample. First, Linear Discriminant Analysis (LDA) was utilized to test whether IFC measurements can separate cellular signatures from five tissue sources: vaginal reference, saliva reference, vaginal cells from hand, and saliva cells from hand, and hand reference. Prediction algorithms based on LDA were then constructed to (1) differentiate vaginal cells from epidermal cells using digital penetration and hand reference samples, and (2) differentiate vaginal cells from saliva cells using digital penetration and oral penetration samples. As part of this study, a framework is proposed for classification of individual cells as originating from either vaginal or epidermal tissue based on probabilities of tissue origin and multivariate distances of individuals cells to each reference group.

## Methods

### Sample collection and preparation

Reference samples of epidermal cells, vaginal cells, and saliva, as well as mock casework oral penetration samples, were collected by study participants pursuant to the Virginia Commonwealth University Institutional Review Board approved protocol #HM20000454_CR7. Reference samples of epidermal cells and vaginal cells, as well as mock casework digital penetration samples, were collected by participants and provided to the Centre for Forensic Science (Toronto & Sault Ste Marie, CA). Written informed consent was obtained from all participants for this study. Epidermal and vaginal reference samples were collected by swabbing the fingers and vaginal cavity respectively for a few seconds using a sterile, pre-wetted cotton swabs (Fisherbrand, P/N 22-363-157 and P/N 22-363-581). For reference saliva samples, ~100μL of saliva was deposited onto a cotton swab. For mock casework samples involving saliva (i.e. oral penetration), participants inserted index and middle fingers into their oral cavity for <30 seconds. Hands were allowed to dry for approximately 10 minutes prior to swabbing as before. For mock casework samples involving digital penetration, samples were collected from participants by self-swabbing fingers at various time points subsequent to vaginal penetration of non-menstruating females. No other constraints were given to participants regarding activity or sample deposition. Following collection, all swabs were allowed to dry and stored at room temperature for up to two months.

Cell populations were eluted from each swab by first incubating in 1.5mL of 1x PBS for 10 minutes followed by vortex for one minute (3000rpm). The resulting cell solution was then filtered through a 100μm mesh to remove swab debris and larger cell aggregates. Next, samples were centrifuged at 1500 × g for 10 minutes at room temperature. The supernatant was removed and the cell pellet was resuspended in 50μL of 1xPBS.

### Imaging Flow Cytometry and statistical analysis

IFC analysis of each cell population (references and mock casework samples) was performed with the Amnis® Imagestream X Mark II instrument (EMD Millipore; Burlington, MA) equipped with 405nm, 488nm, 561nm, and 642nm lasers. Laser voltages for all tests were set at 120mW, 100mW, 100mW and 150mW, respectively. Images of individual events were captured in five detector channels labeled: 1 (430-505nm), 2 (505-560nm), 3 (560-595nm), 5 (640-745nm), and 6 (745-780nm). Channel 4 was used to capture brightfield images. Magnification was set at 40x and autofocus was enabled so that the focus varied with cell size. Examples of cell images collected across multiple wavelengths are provided in Figure 1.

**Figure 1.**
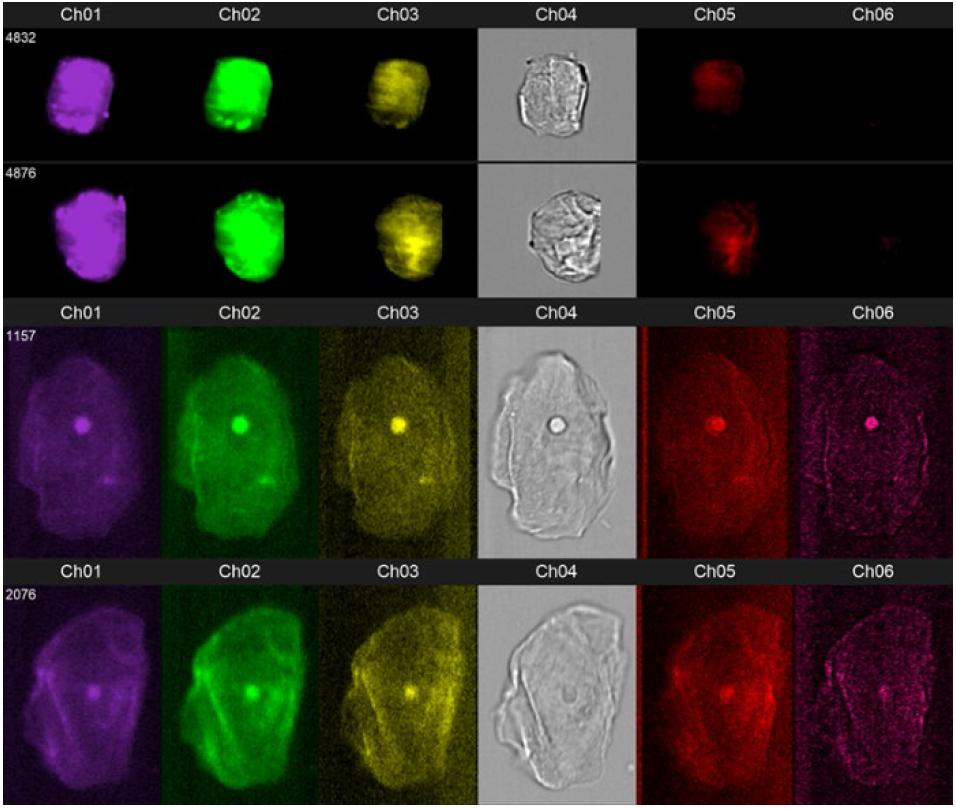
IFC images collected from six detector channels for two epidermal cells (top two rows) and two vaginal cells (bottom two rows). The horizontal axis in each image is 80μm.

Because IFC captures data for all objects ranging in diameters between ~1 μm and ~150μm which can include a range of both cellular and non-cellular material (e.g., biological debris, larger cellular aggregates and/or non-biological particles), the population of objects corresponding to individual, intact cells had to be identified in each sample first. To accomplish this, raw image files (.rif) generated from IFC were imported into IDEAS® Software (EMD Millipore; Burlington, MA). Individual cells were differentiated from debris or other non-cellular material by filtering for objects with calculated areas between 1000-2500 μm^2^ (determined from the ‘Area_Ch04’ variable) and Gradient RMS Ch04 values greater than 50 which corresponds to cells that are in focus within the brightfield detector channel.

To enhance the selectivity for vaginal tissue, a subpopulation of cells was identified that was abundant in vaginal reference swabs and relatively scarce in hand reference swabs. This was defined by cell area in the brightfield channel (>1000 μm^2^) and contrast value in the brightfield channel (<10) (Table S1). The subpopulation was then extracted from each tissue sample in this study used for both signature identification and classification tests.

Cellular measurements were extracted in the Ideas software. Seven different measurement categories were used; area, aspect ratio, contrast, intensity, mean pixel, brightness detail intensity at ‘R3’ pixel increment, brightness detail intensity at ‘R7’ pixel increment, compactness, and the ratio of intensity values. Within each of these categories, data was extracted from each of the six detector channels yielding a total of 87 measurements for each cell.

The resulting dataset was exported to SPSS v28 (IBM, Inc. Chicago, IL) for statistical analysis. Multivariate modeling and classification tests among cell populations were based on Linear Discriminant Analysis (LDA) using a within-group covariance matrix. Distance calculations involved in these functions utilized the squared Mahalanobis Distance described in [4]. Probabilities of group membership were calculated using the formula below where Dj represents the covariance matrix for discriminant functions of the jth group and X^2^ represents the squared Mahalanobis Distance to the same reference group’s centroid/multivariate mean.

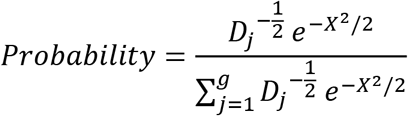

For this study, classification functions were generated for two different applications: (1) differentiating non-epidermal cells (e.g., vaginal or saliva) from epidermal cells and (2) differentiating vaginal cells from saliva cells. The overall scheme workflow for classifying cells from an unknown/blinded sample are shown in Figure S1.

### DNA extraction and quantification of mock casework digital penetration samples

Samples were subjected to differential extraction to isolate DNA. First, 505μl of epithelial extraction buffer ([10mM Tris-HCl (pH 8.0), 100mM sodium chloride, 1mM EDTA (pH 8.0), (TNE)], 1% SDS, and 0.2 mg/ml proteinase K) was added to samples previously prepared in 2 ml tubes. These were then incubated on a rocking platform at 37 °C for 2 h. Once incubation was complete, samples were placed in spin baskets and subjected to centrifugation at 18,000 xg for five minutes. The swab was then placed in a separate tube for storage at −20 °C. The liquid [epithelial fraction] was placed in a new tube for purification. The remaining pellet [equivalent to the sperm fraction if being processed for casework] was subjected to a series of three washes (one with TNE and two with dH2O). Following the removal of the final supernatant, 400 μl of sperm extraction buffer (10mM Tris-HCl (pH 8.0), 100mM sodium chloride, 1mM EDTA (pH 8.0), 2.5% sarkosyl, 0.39M dithiothreitol, and 0.5 mg/ml proteinase K) was added to each tube, and the tubes were incubated on a rocking platform at 37 °C for a minimum of 2 h. Following incubation, sperm fractions were transferred into new 2 ml screw top vials for purification. Epithelial and sperm fractions were both subjected to purification using the Qiagen EZ1® Advanced XL system, buffer MTL and the large volume protocol with elution in 40 μl TE, as per manufacturer’s guidelines.

Aliquots (2 μl) were removed from the eluted samples post-purification and quantified using quantitative Real Time polymerase chain reaction using the Bio-Rad CFX 96™ Real-Time PCR Detection System (Bio-Rad, Hercules, CA) and Promega Plexor®HY Quantitation System (Promega Corporation, Madison, WI).

## Results and Discussion

### Isolating and refining a vaginal signal distinguishable from other epithelial cell types

#### 1. Comparison of IFC measurements across reference cell types

Initial IFC comparisons of vaginal and epidermal cell populations from reference samples indicated clear differences in cell size (average area <900 μm^2^ for epidermal, >1000 μm^2^ for vaginal cells) as well as optical contrast values in the brightfield channel which are primarily a function of cell thickness. These attributes were used to define a subpopulation of cells that were abundant in vaginal swabs and relatively scarce in hand swabs (Table S1 and described in Methods). Within the resulting subpopulation of large and low contrast cells, features consistent with nuclei were observed in vaginal cell populations but were absent in hand epidermal reference samples. Cells from reference hand swabs that met criteria for this subpopulation were a mixture of single cells and, occasionally, clusters of two or more cells. However, these instances of individual cells still did not exhibit obvious nuclei. Importantly, these observations are consistent with well-established histologic differences between vaginal and epidermal tissue [5,6] as well as previously described morphological differences between vaginal and epidermal cells recovered from aged biological samples [2].

#### 2. Multivariate modeling of IFC differences across references and mock casework samples

Next, discriminant analysis was used to model differences between cell populations derived from reference samples and fingers swabbed subsequent to digital or oral penetration using the full set of IFC measurements. To enhance the relevance for forensic casework, we first tested whether cell populations derived from reference samples were distinguishable from those derived from the same tissue deposited under mock casework conditions, as described above. Figure 2 (left panel) shows the separation of cell populations derived from (1) swabs of vaginal cavity, (2) swabs of the oral cavity, (3) swabs of fingers following oral penetration, and (4) swabs of fingers following digital penetration along the first two multivariate discriminant functions. Spatially resolved clustering is observed among all four cell populations with distances between group centroids showing significant differences (MANOVA/Wilk’s Lambda= 0.28, p<0.0001). Interestingly, the largest multivariate distances were observed between cell populations from vaginal swabs and cell populations from vaginal fluids deposited onto fingers (blue vs. orange clusters). Similarly, cell populations derived from saliva reference samples showed distinct clustering along discriminant functions compared to saliva that was dried onto and then sampled from fingers (red versus green clusters).

**Figure 2.**
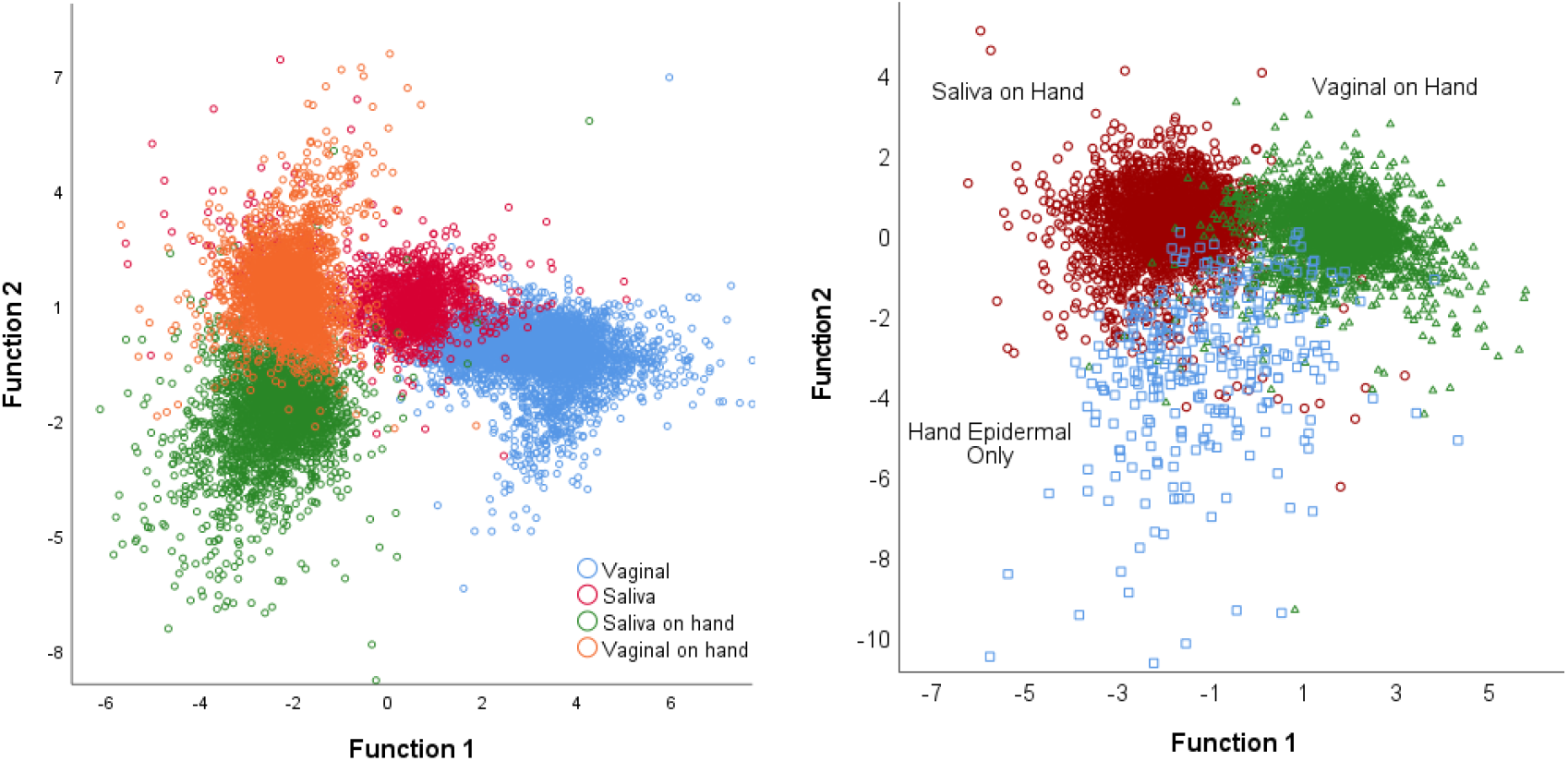
Multivariate comparisons of cell populations. Left plot shows LDA of cell populations from four sources: vaginal swab, saliva, finger swab subsequent to oral penetration (saliva on hand), and finger swab subsequent to digital penetration (vaginal on hand). Right plot shows LDA of cell populations from: finger swab (hand epidermal only), finger swab after oral penetration (saliva on hand), and finger swab after digital penetration. Each data point represents a single cell. Each axis is a linear combination of original IFC measurements.

The fact that non-epidermal epithelial cells collected from fingers subsequent to vaginal or oral contact differ from those collected directly from source tissue is not unexpected. Previous studies have reported differences in the abundance and diversity of epithelial cell types recovered from buccal swabs compared to those found within saliva samples as well as in the quantity and quality of DNA [7,8]. This variation was attributed in part to swab samples having epithelial cells from deeper tissue layers compared to fluid samples predominantly composed of cells shed from the outer tissue layers [8]. In this study, review of the IFC images showed no obvious visual differences between the four cell populations from vaginal tissue and saliva (saliva, oral penetration, vaginal, digital penetration). However, variable loadings from the first two discriminant functions indicate that fluorescence intensity in channels 2 (505-560nm) and 3 (560-595nm) are important factors in the discriminant functions from Figure 2 which may reflect differences in biochemical profiles and/or proportion of epithelial cell types between samples from swab collection versus vaginal/saliva fluid.

Based on these results, additional multivariate discriminant analyses explored the differences in cell populations derived from swabbed fingers subsequent to different activities. A comparison of cell populations from fingers swabbed subsequent to (1) vaginal or (2) oral contact, and (3) without such contact, showed strong multivariate differences, Wilks lambda =0.013, p<0.001. Each cell population type showed well-resolved clustering with minimal overlap (Figure 2 right panel), indicating robust multivariate differences in IFC measurements.

### Developing and testing algorithms for classifying vaginal and epidermal cells

From the multivariate differences observed in Figure 2, LDA algorithms for classifying cells as originating from either epidermal or vaginal tissue were constructed using IFC measurements as primary variables. Two separate classification procedures were conducted: first, comparing samples from digital penetration to hand reference samples, and second, comparing samples from digital penetration to oral penetration. In both cases, the posterior probability of each cell belonging to one of the tissue sources was calculated. For the first test comparing cell populations from digital penetration and from hand references, the LDA model was trained on all the sample cell populations shown in Table 1 except for one sample cell population which was held-out and tested in a blind fashion with posterior probabilities calculated for each cell. This process was repeated with different sample cell populations held-out and tested blindly for every iteration. The second test comparing cell populations from digital penetration to cell populations from oral penetration was conducted similarly. In each case, the held-out test population refers to the entire sample of cells collected from a single human participant subsequent to swabbing.

**Table 1.** Cell Populations from Digital Penetration Mixtures and Hand Reference Samples.

| **Sample ID** | **Cell Population Source** | **Time Before Collection, Activity<sup>1</sup>** | **# Cells DigPen<sup>2</sup>** | **Cells > .99PP** | **Cells > .95PP** | **Mean PP** | **Mean Distance** |
| --- | --- | --- | --- | --- | --- | --- | --- |
| **14A** | Dig. Pen. | <15min | 407<br>(451) | 167 | 290 | 0.94 | 1.97 |
| **16A** | Dig. Pen. | <15min | 373<br>(410) | 101 | 226 | 0.94 | 1.54 |
| **19B** | Dig. Pen. | <15min | 258<br>(262) | 243 | 252 | 0.96 | 0.47 |
| **21B** | Dig. Pen. | 20hrs,<br>multiple<br>washes | 9 (18) | 2 | 7 | 0.97 | 1.46 |
| **23B** | Dig. Pen. | <15min | 1381<br>(1389) | 1196 | 1351 | 0.99 | 0.26 |
| **23B** | Dig. Pen. | 3 hours | 151<br>(192) | 15 | 53 | 0.86 | 2.89 |
| **30B** | Dig. Pen. | <15min | 222<br>(222) | 219 | 220 | 1.00 | 5.01 |
| **1** | Hand Ref. |  | 4 (28) | 1 | 2 | 0.91 | 1.90 |
| **5** | Hand Ref. |  | 20 (39) | 11 | 13 | 0.92 | 1.66 |
| **6** | Hand Ref. |  | 3 (43) | 1 | 1 | 0.82 | 2.44 |
| **8** | Hand Ref. |  | 1 (27) | 0 | 0 | 0.75 | 3.35 |
| **9** | Hand Ref. |  | 9 (32) | 4 | 4 | 0.91 | 1.92 |
| **11** | Hand Ref. |  | 2 (28) | 1 | 1 | 0.76 | 64.56 |
| **19B** | Hand Ref. |  | 4 (6) | 3 | 4 | 0.99 | 1.90 |
| **20A** | Hand Ref. |  | 4 (5) | 3 | 3 | 0.96 | 1.09 |
| **23B** | Hand Ref. |  | 43 (75) | 19 | 24 | 0.89 | 2.58 |
| **A24** | Hand Ref. |  | 0 | 0 | 0 | N/A | N/A |
| **D61** | Hand Ref. |  | 71 | 11 | 16 | 0.77 | 15.27 |
| **P94** | Hand Ref. |  | 2 (436*) | 0 | 0 | 0.63 | 3.75 |

<sup>1</sup>Unless otherwise indicated, no hand washing took place prior to sampling
<sup>2</sup>Number of cells initially classified as epithelial cells from digital penetration. The total number of cells detected in each sample shown in parentheses.

Results from classification tests of digital penetration samples compared to hand reference samples are shown in Table 1. Samples that were collected less than 15 minutes after digital penetration showed relatively large numbers of cells—ranging from a low of 222 cells to a high of 1381—classifying as vaginal, *i.e.* possessing a posterior probability of vaginal group membership higher than 0.50. The average posterior probability for these cells ranged between 0.94 and 1.00. Samples with longer periods of time between digital penetration and swabbing showed comparatively lower numbers of cells classifying as vaginal, *e.g.,* 151 cells for sample 23B (3 hours post activity) and 9 cells for sample 21B (20 hours post activity with multiple washes), with average posterior probabilities of 0.86 and 0.97, respectively.

In comparison, samples collected from fingers without any known contact with body fluids showed fewer cells classifying as vaginal, ranging from a low of 0 cells to a high of 71, with fewer than 10 cells classifying as vaginal in the majority of samples (Table 1). For the one sample possessing more than fifty cells classifying as vaginal, the average posterior probability (0.77, sample D61) was low relative to digital penetration samples. Given the variability of both the number of cells classifying as vaginal in these samples as well as posterior probability, there are several possible sources for these cells. First, some percentage of these cells may in fact be vaginal cells that were transferred directly or indirectly (*i.e*., through common living spaces and activities of the participants) and persist on the individual’s hands at the time of collection. A second possibility is that these are cells from other layers of the epidermis which may have different physical and/or chemical properties from more surficial corneocytes that comprise that majority of biological material in touch/trace samples [9,10]. Third, these cells may represent contributions from other epithelial cell sources such as mucosal epithelium or saliva that are transferred in small quantities to an individual’s hands and are subsequently misclassified as vaginal cells (discussed further below). Although there is no clear-cut way to distinguish between these possibilities in this study, future work could explore coupling IFC imaging with fluorescent probes targeting tissue-specific protein and/or genetic markers within the cell to differentiate cell sources (e.g., micro-RNA [11], cytokeratin alleles [12].

### DNA profiling of digital penetration samples

DNA from each of the digital penetration samples was extracted, quantitated, and typed. Results showed that each of the five samples collected less than 15 minutes after digital vaginal penetration showed high DNA yields (between 20ng-788ng) with DNA profiles that typed as single source female (Table 2). Sample 23B-2, which was collected ~3 hours after activity, also showed high DNA yields (~44ng) and a mixed DNA profile that was 20:1 female:male. Unsurprisingly, sample 21B, which was collected >20 hours post digital penetration and after multiple hand washes, showed low quantities of total DNA (0.71ng) and a DNA profile that was single source male. Overall, the DNA findings are consistent with expectations in light of cell classification results, *i.e.* samples with high numbers of cells classifying as vaginal correspondingly show high DNA yields and single source female profiles, and the one sample with few cells classifying as vaginal yielded much less DNA, with an undetectable female profile.

**Table 2.** DNA Quantification from Mock Digital Penetration Samples.

| **Sample ID** | **Time & Activity<sup>1</sup>** | **[Auto] pg/μl** | **[Y] pg/μl** | **[Auto]/[Y]** | **Total DNA (ng)** | **DNA Interpretation** |
| --- | --- | --- | --- | --- | --- | --- |
| **14A** | <15min | 19350 | 2.22 | 8707 | 735.30 | single source female |
| **16A** | <15min | 758.80 | N/A | N/A | 28.83 | single source female |
| **19B** | <15min | 19380.0 | 0.91 | N/A | 736.44 | single source female |
| **21B** | 20hrs, multiple washes | 18.73 | 5.59 | 3.4 | 0.71 | single source male |
| **23B-1** | <15min | 11320 | 10.14 | 1117.0 | 430.16 | single source female |
| **23B-2** | 3 hours | 1165.0 | 22.33 | 52.2 | 44.27 | ~20:1 (female:male) |
| **30B** | <15min | 20750 | 0.24 | 86040 | 788.50 | single source female |

<sup>1</sup>Unless otherwise indicated, no hand washing took place prior to sampling

### Differentiating vaginal cells from saliva

Within a casework context, saliva may be an alternative explanation for recovering a complainant’s DNA from a suspect’s hands where digital vaginal penetration is alleged. Therefore, the second classification test focused on resolving vaginal cell populations from saliva cell populations by comparing oral penetration samples (*i.e.* swabs of fingers that had been inserted in oral cavity) to digital penetration samples. Results showed oral penetration samples contained a relatively small proportion of cells with posterior probabilities to vaginal tissue greater than 50%, ranging from 0 cells (P94) to 99 cells (K47), which constituted between 0 and ~12% of the total cell population (Table 3). These cells had lower average posterior probabilities (0.74-0.88) for vaginal tissue compared to digital penetration samples. The higher numbers of cells in saliva samples that exhibit morphological and/or autofluorescence properties similar to vaginal tissue was unsurprising given the morphological, structural, and biochemical similarities between these tissue types [5]. Additionally, previous IFC characterizations have observed smaller variation between saliva and vaginal cells as compared to epidermal tissue [2].

**Table 3.** Cell Populations from Digital Penetration Mixtures and Saliva Hand Mixtures.

| **Sample ID** | **Cell Population Source** | **Time before Collection, Activity<sup>1</sup>** | **# Cells DigPen<sup>2</sup>** | **# Cells Saliva<sup>3</sup>** | **Cells > .99PP** | **Cells > .95PP** | **Mean PP** | **Mean Dist.** |
| --- | --- | --- | --- | --- | --- | --- | --- | --- |
| **14A** | Dig. Pen. | <15min | 252 | 155 | 18 | 47 | 0.82 | 3.547 |
| **16A** | Dig. Pen. | <15min | 21 | 352 | 2 | 4 | 0.78 | 2.951 |
| **19B** | Dig. Pen. | <15min | 258 | 0 | 247 | 256 | 0.99 | 0.523 |
| **21B** | Dig. Pen. | 20hrs, multiple washes | 9 | 0 | 6 | 8 | 0.98 | 0.585 |
| **23B** | Dig. Pen. | <15min | 1353 | 28 | 737 | 1104 | 0.96 | 0.534 |
| **23B** | Dig. Pen. | 3 hours | 140 | 11 | 31 | 69 | 0.90 | 1.811 |
| **30B** | Dig. Pen. | <15min | 221 | 0 | 221 | 221 | 1.00 | 22.986 |
| **A24** | Saliva on Hand | <15min | 62 | 640 | 6 | 14 | 0.77 | 2.611 |
| **D61** | Saliva on Hand | <15min | 17 | 527 | 5 | 6 | 0.82 | 4.625 |
| **I66** | Saliva on Hand | <15min | 20 | 188 | 10 | 13 | 0.88 | 3.609 |
| **K47** | Saliva on Hand | <15min | 99 | 789 | 6 | 13 | 0.74 | 3.964 |
| **P94** | Saliva on Hand | <15min | 0 | 123 | 0 | 0 | N/A | N/A |

<sup>1</sup>Unless otherwise indicated, no hand washing took place prior to sampling
<sup>2</sup>Number of cells initially classified with digital penetration reference group of epithelial cells with posterior probabilities >50%.
<sup>3</sup>Number of cells initially classified with saliva reference group of epithelial cells with posterior probabilities >50%.

When digital penetration samples were classified as either vaginal or saliva using the same algorithms, cell populations from four of the samples (19B, 23B-1, 23B-2, and 30B) were overwhelmingly identified as vaginal cells (92-100% of the cell population; Table 3). Cell populations from these samples also exhibited high mean posterior probabilities ranging between 0.90 and ~1.00. However, two digital penetration samples showed a large subset of cells similar to saliva: in sample 14A, 155 cells (approximately 38% of the cell population), and in sample 16A, 352 cells (approximately 94% of the cell population). Within these two samples, the subset of cells classifying as vaginal were also associated with lower average posterior probabilities; 0.82 and 0.78, respectively.

For these two samples, the higher proportion of cells with posterior probabilities greater than 50% for saliva may represent the true presence of saliva (since samples were created and collected by individual donors and donation did not preclude using saliva as a lubricant). Alternatively, these cells may have originated from vaginal tissue but were misclassified as saliva due to a level of variation either between tissue groups or within tissue groups; variation which is not adequately captured by the reference dataset used in this study. If the latter is true, classification algorithms may be improved by sampling additional donors and/or generating samples with a greater variety of depositional circumstances.

Because of the apparent heterogeneity in cell populations from samples 14A and 16A (and the potential for contributions of multiple tissue types to these samples), we tested whether classification of the remaining digital penetration samples could be improved by removing 14A/16A from the reference dataset. Results showed different trends across the remaining samples. For example, the average posterior probability for 23B (cells collected <15min after activity) increased slightly from ~0.96 to 0.99. Conversely, probability decreased for the second 23B sample (cells collected ~ 3 hours after activity) from 0.90 to 0.79 and 19B was largely unchanged (0.99 for each classification). The varying results in classification indicate that exclusion of 14A/16A does not have a systematic effect on tissue classification which may in part be a function of the number of reference donor cell populations that remain after removal and/or that predictive functions are heavily influenced by other donor cell populations within this group.

## Conclusions

Taken together, these results indicate that cell populations recovered from hand swabs following digital penetration yield greater numbers of cells consistent with vaginal tissue and higher average posterior probabilities for vaginal tissue compared to cell populations recovered from hand swabs without any prior activity. Within the context of casework, these differences could be used to create minimum thresholds by which vaginal tissue is initially inferred from hand swabs. For example, based on this dataset requiring that an unknown sample yield more than 50 cells classifying as vaginal, with a mean posterior probability of at least 0.85 would eliminate all false positive detection of vaginal cells in both hand reference samples and licked fingers. While one of the digital penetration samples did not meet this threshold (21B which showed 9 cells classifying as vaginal tissue with a mean posterior probability of 0.97), this sample was collected 20 hours after digital penetration, with several intervening hand washings. It is possible that there were very few vaginal cells remaining on the subject’s hands at the time of collection. In any case, any interpretation threshold will have some effect on sensitivity, particularly on samples at the edge of detectability.

Importantly, each of the digital penetration samples that showed higher numbers of cells classifying as vaginal tissue as well as higher posterior probabilities for vaginal tissue also yielded quantities of DNA consistent with the presence of large quantities of nucleated epithelial cells. Additionally, these samples produced DNA profiles that were either single source female or majority female (20:1) as would be expected following the deposition of vaginal material. This suggests that the cellular signatures characterized with IFC may be associated with the presence and/or abundance of DNA originating from vaginal tissue. The detection of vaginal cells in the sample collected three hours following digital penetration indicates the potential persistence of vaginal cells for longer, more relevant periods of time for law enforcement following the suspected activity. Further, the proposed vaginal cell signatures may be probative when saliva is not a proposed mechanism for DNA deposition, e.g., cases of object penetration or instances when the detection of vaginal fluid on clothing or other substrates may be probative. Future work should continue to examine how vaginal cell signatures change with increasing time-since-depositions as well as the effect that various post-depositional activities have on persistence of signatures (e.g., hand-washing or wiping).

Differentiation of vaginal cell populations from saliva cell populations was also promising in that saliva deposits showed markedly lower probabilities of a vaginal tissue source compared to the majority of digital penetration samples in this study. These differences suggest that thresholds for cell count and/or average posterior probability for inferring the presence of vaginal cells when saliva is a possible component of the sample. From this dataset, a threshold of 50 cells and an average posterior probability of 0.80 would eliminate false positive identification of vaginal cells within reference mixtures of saliva and epidermal cells. With this same threshold, vaginal tissue would not be detected in two of the seven digital penetration samples (i.e., false negative).

While these results indicate that morphological and autofluorescence properties of individual cells can discriminate between saliva, epidermal, and vaginal tissue sources in forensically relevant mixture samples, future efforts should continue characterize cell populations in comparable mixture samples that also have a range of time-since-depositions and post-depositional activities. This will help establish the sensitivity and/or limit of detection for this approach, e.g., the minimum number of cells and/or posterior probability required to infer the presence of vaginal tissue or the maximum time-since-deposition of a sample in which vaginal cells may be detected. These thresholds could then be set to optimize relative rates of false positive/false negative identifications and place its sensitivity within the context of other long-standing serological techniques.

## Acknowledgements

This project was funded by the Center for Innovative Technology-Commonwealth Research Commercialization Fund (MF-19-013-LS; PI Ehrhardt). Flow cytometry services in support of the project were provided by the VCU Massey Cancer Center, supported in part with funding from NIH-NCI P30CA016059. The sponsoring agencies were not involved in the study design; collection, analysis and interpretation of data, or the decision to submit the article for publication.

## References

[1] A.B. Hall, R. Saferstein, Forensic Science Handbook, CRC Press, Third edition. | Boca Raton, FL: CRC Press, 2019-,2020. https://doi.org/10.4324/9781315119939.

[2] E.R. Brocato, M.K. Philpott, C.C. Connon, C.J. Ehrhardt, Rapid differentiation of epithelial cell types in aged biological samples using autofluorescence and morphological signatures, PLoS One. 13 (2018) e0197701. https://doi.org/10.1371/journal.pone.0197701.

[3] S. Ingram, A. DeCorte, M.K. Philpott, T. Moldenhauer, S. Stadler, C. Steinberg, J. Millman, C.J. Ehrhardt, Differentiation of vaginal cells from epidermal cells using morphological and autofluorescence properties: Implications for sexual assault casework involving digital penetration, Forensic Sci Int Genet Suppl Ser. 8 (2022) 17–19. https://doi.org/10.1016/j.fsigss.2022.09.007.

[4] Morrison DF, Multivariate statistical methods (McGraw-Hill series in probability and statistics), 2nd ed., McGraw-Hill, New York, 1976.

[5] H. Evers, C.G. Birngruber, F. Ramsthaler, U. Müller, S. Brück, M.A. Verhoff, [Differentiation of epithelial cell types by cell diameter]., Arch Kriminol. 228 (2011) 11–9.

[6] G. Plewig, R.R. Marples, Regional Differences of Cell Sizes in the Human Stratum Corneum. Part I, Journal of Investigative Dermatology. 54 (1970) 13–18. https://doi.org/10.1111/1523-1747.ep12551482.

[7] N.L. Rogers, S.A. Cole, H.-C. Lan, A. Crossa, E.W. Demerath, New saliva DNA collection method compared to buccal cell collection techniques for epidemiological studies, American Journal of Human Biology. 19 (2007) 319–326. https://doi.org/10.1002/ajhb.20586.

[8] C. Theda, S.H. Hwang, A. Czajko, Y.J. Loke, P. Leong, J.M. Craig, Quantitation of the cellular content of saliva and buccal swab samples, Sci Rep. 8 (2018) 6944. https://doi.org/10.1038/s41598-018-25311-0.

[9] C.E. Stanciu, M.K. Philpott, Y.J. Kwon, E.E. Bustamante, C.J. Ehrhardt, Optical characterization of epidermal cells and their relationship to DNA recovery from touch samples, F1000Res. (2015). https://doi.org/10.12688/f1000research.7385.1.

[10] J. Burrill, B. Daniel, N. Frascione, A review of trace “Touch DNA” deposits: Variability factors and an exploration of cellular composition, Forensic Sci Int Genet. 39 (2019). https://doi.org/10.1016/j.fsigen.2018.11.019.

[11] S. Seashols-Williams, C. Lewis, C. Calloway, N. Peace, A. Harrison, C. Hayes-Nash, S. Fleming, Q. Wu, Z.E. Zehner, High-throughput miRNA sequencing and identification of biomarkers for forensically relevant biological fluids., Electrophoresis. 37 (2016) 2780–2788. https://doi.org/10.1002/elps.201600258.

[12] M. Katherine Philpott, C.E. Stanciu, Y.J. Kwon, E.E. Bustamante, S.A. Greenspoon, C.J. Ehrhardt, Analysis of cellular autofluorescence in touch samples by flow cytometry: implications for front end separation of trace mixture evidence, Anal Bioanal Chem. 409 (2017) 4167–4179. https://doi.org/10.1007/s00216-017-0364-0.

