## Supplemental Table and Figure for "Differentiation of vaginal cells from epidermal cells using morphological and autofluorescence properties: Implications for sexual assault casework involving digital penetration"

**Table S1. Cell Subpopulations in Reference Vaginal and Epidermal Samples**

| Reference Tissue* | Cell Count: Contrast <10 | Cell Count: Contrast <10 and Area >1000 $\mu m^3$ |
|---|---|---|
| **Vaginal (23A)** | 695 | 555 |
| **Vaginal (23A)** | 648 | 547 |
| **Vaginal (23A)** | 515 | 404 |
| **Vaginal (20B)** | 191 | 123 |
| **Vaginal (11)** | 642 | 549 |
| **Vaginal (11)** | 678 | 570 |
| **Vaginal (10)** | 453 | 231 |
| **Hand (1)** | 28 | 8 |
| **Hand (5)** | 39 | 10 |
| **Hand (6)** | 43 | 10 |
| **Hand (8)** | 27 | 4 |
| **Hand (9)** | 32 | 6 |
| **Hand (11)** | 28 | 6 |
| **Hand (D61)** | 19 | 1 |
| **Hand (20A)** | 5 | 0 |
| **Hand (23B)** | 75 | 9 |
| **Hand (19B)** | 6 | 0 |

***Sample ID shown in parentheses**

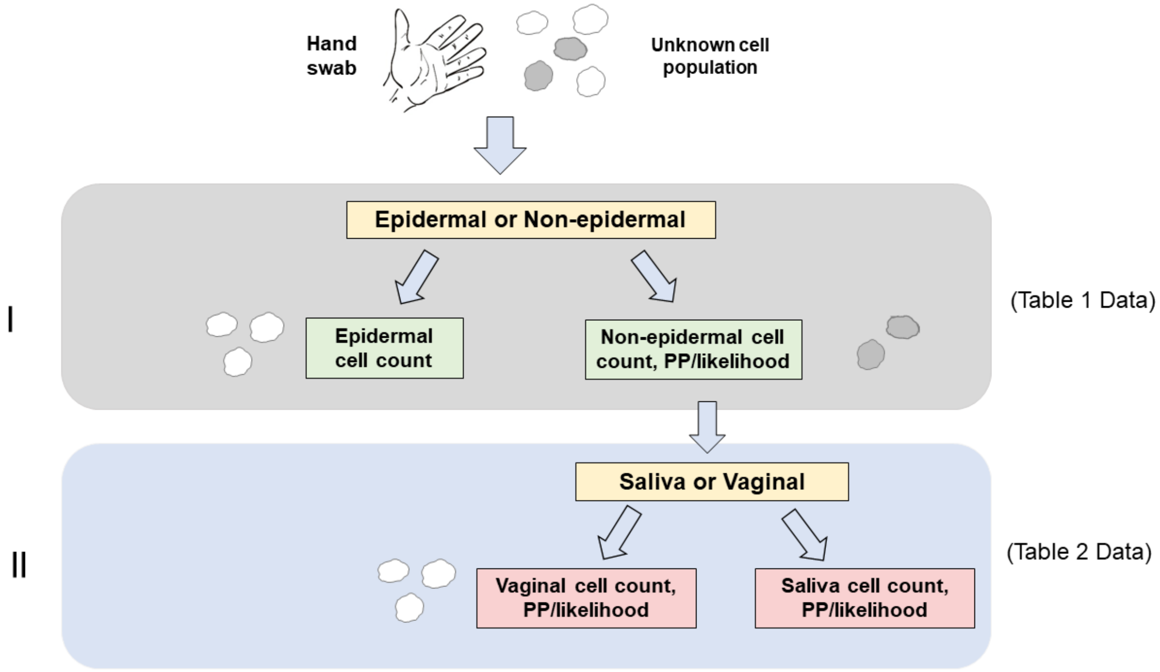


**Figure S1**. Conceptual flow chart for classification of cell populations from an unknown hand swab sample
